# Characterization of the Complete Chloroplast Genome Sequences and Phylogenetic Relationships of Four Oil-Seed Camellia spp. and related taxa

**DOI:** 10.1101/2023.10.03.560681

**Authors:** Huihua Luo, Boyong Liao, Yongjuan Li, Runsheng Huang, Kunchang Zhang, Longyuan Wang, Jing Tan, Yuzhou Lv, Can Lai, Yongquan Li

## Abstract

Some species in the Sect. *Oleifera* of the genus *Camellia* L. known as oil-seed camellia because of their high oil content and economic value. Additional studies aimed at clarifying the phylogenetic relationships and chloroplast genomes of *Camellia* species are needed to hybridization, as well as improve the breeding, selection and interspecific hybridization of *Camellia* species. The complete chloroplast genomes (cpDNA) of the four oil-seed camellia species *C. semiserrata, C. meiocarpa, C. suaveolens*, and *C. osmantha* were resequenced to clarify their interspecific relationships. These cpDNA had typical tetrad structures, and they were highly conserved in various structural features. The total lengths of the cpDNA ranged from 156,965 to 157,018 bp, and 134 genes were annotated, including 88 protein-coding genes, 37 transfer RNA genes, and 8 messenger RNA genes. The average GC content of these genomes was 37.3%. The codons with the highest and lowest codon usage bias were UUA (which codes for leucine) and AGC (which codes for serine), respectively. The number of simple sequences repeats of the four *Camelia* species ranged from 38 to 40. Mononucleotide repeats were the most common repeat type, followed by tetranucleotide, trinucleotide, and hexanucleotide repeats. Our phylogenetic analysis of cpDNA, coupled with the results of previous ploidy analyses and artificial interspecific hybridization, revealed that *C. semiserrata* was most closely related to *C. azalea, C. suaveolens* was most closely related to *C. gauchowensis, C. osmantha* was most closely related to *C. vietnamensis*, and *C. meiocarpa* was most closely related to *C. oleifera*. The phylogenetic relationships between oil-seed camellia species with high oil content and economic value were characterized. Our analysis of the cpDNA provided new insights that will aid the use of artificial distant hybridization in camellia breeding programs.

## Introduction

Oil-seed camellia plants have become economically important woody oil plants in China in recent years. Some plants in the genus *Camellia* L. (family Theaceae) are generally referred to as oil-seed camellia because of their high oil content and economic value (FRPS,1998; State owned forest farm and forest seedling work station of the State Forestry Administration, 2016). Oil-seed camellia plants are unique woody oil plants and the fourth largest in the world in terms of production after *Olea europaea* L., *Elaeis guineensis* Jacq., and *Cocos nucifera* L.. Camellia oil is highly nutritious, and its unsaturated fatty acid content is up to 90% higher than that of other common edible oils on the market. Oil from camellia seeds has been shown to be effective in preventing cardiovascular and cerebrovascular diseases. Camellia oil is also considered a healthy edible vegetable oil by the Food and Agriculture Organization. It has various uses in industry, such as the production of cosmetic products and medicine. In addition, the potential applications of these residues from the oil pressing process are manifold, and additional studies will likely broaden the uses of these residues (Li et al., 2011).

Oil-seed camellia plants have been cultivated for approximately 2, 300 years (Wang et al., 2020). China contributes approximately 95% of the world’s production of *Camellia* plants and 90% of the world’s production of *Camellia* seeds. The largest oil-tea cultivating areas in China are in the following provinces of Hunan, Jiangxi, Guangxi, Zhejiang, Fujian, and Guangdong. Oil-seed camellia is also grown in 1, 537 counties (cities) in 14 other provinces in China. The main oil-seed camellia tree species cultivated in China are *C. oleifera, C. meiocarpa, C. gauchowensis*, and *C. semiserrata*. Edible oil is the main product of *Camellia* cultivation.

Although various classifications of the genus *Camellia* have been proposed, these classifications have mostly been based on morphological characters and molecular information from DNA sequences and chloroplast sequences (Chang, 1981). Given that hybridization and polyploidization are frequent in this group, traditionally used morphological indexes are likely affected by various environmental factors such as topography, soil, and climate. Trees reconstructed based on a few genes and chloroplasts often show inconsistent or reticulate evolution. This poses a major challenge to taxonomic and phylogenetic analyses of the genus *Camellia* L. and has greatly limited progress in our understanding of the classification of *Camellia* L. (Min et al., 1996; Hong, 1981). The phylogenetic relationships among *Camellia* members remain unclear. Other non-morphological sources of data are needed to clarify their evolutionary relationships, such as, whole genome information, chromosome variation and the utility of artificial interspecific hybridization (Zhang et al., 2022; Yu et al., 2022; Ye et al., 2021; Zhong et al., 2020; Chang et al., 2016; Zhou et al., 2001).

The chloroplast is an important site for energy conversion and photosynthesis in green plants. Chloroplast genomes (cpDNA) are one of the three major genetic systems in plants. They have been widely used in evolutionary studies because of their low nucleotide substitution rates, uniparental inheritance, conserved structure, and low molecular weight (Tang et al., 2022; Tian et al., 2021; Zhu et al., 2022). The cpDNA of these species have been resequenced several times to clarify their relationships, such as the newly described oil-seed species of *C. osmantha* (Ma et al., 2012; Liu et al., 2021). Here, we used next-generation high-throughput sequencing to assemble, annotate, and characterize the cpDNA of four *Camellia* species. We generated more chloroplast genomic resources for oil-seed *Camellia* to characterize the structure of cpDNA and clarify the relationships among these four oil-seed camellia species within the genus *Camellia* L. Our findings provide new insights into chromosome variation and the utility of artificial interspecific hybridization for camellia breeding. The results of our study will also aid future studies aimed at genetically improving the oil production of several oil-seed camellia species.

## Materials And Methods

### Experimental Materials and Sequencing

Seeds of *C. semiserrata, C. meiocarpa, C. suaveolens*, and *C. osmantha* were collected from Zhaoqin, Meizhou, Lechang, and Nanning in southern China in 2013 (**Table 1**). The seedlings of four camellia species were planted in Xiaokeng state-owned forest farm (24°15’ N, 113°35’ E) in Qujiang District, Shaoguan City, Guangdong Province, China in 2015. Young leaves of four species without signs of pests and disease were collected in bags filled with dried silica gel prior to transport to the laboratory in 2020. All four species were identified by the Sun Yat-sen University Herbarium. The remaining silica gel-dried young leaves and vouchers were deposited in the Zhongkai University of Agriculture and Engineering Herbarium for subsequent studies.

**Table 1.** Sample sequencing data information.

| Species | Voucher NO. | Seed sample collection sites | Seed collection year | Raw base (BP) | Raw Q30(%) |
| --- | --- | --- | --- | --- | --- |
| *C. semiserrata* Chi. | ZKU2020032 | Guangning District, Zhaoqin City, Guangdong Province, China | 2013 | 11, 788, 927, 200 | 90.56;<br>87.11 |
| *C. meiocarpa* Hu. | ZKU2020035 | Xingning District, Meizhou City, Guangdong Province, China | 2013 | 9, 157, 311, 900 | 91.22;<br>86.76 |
| *C. suaveolens* Ye. | ZKU2020051 | Lechang City, Guangdong Province, China | 2013 | 8, 529, 454, 200 | 91.67;<br>88.99 |
| *C. osmantha* Ye. | ZKU2020159 | Guangxi Forestry Research Institute, Nanning, China | 2013 | 10, 930, 712, 700 | 91.53;<br>87.79 |

A modified CTAB method was used to extract total DNA (Rogers & Bendich, 1989). The Illumina NovaSeq platform was used to construct, quality, and paired-end sequence (2×150bp) the DNA libraries using the Illumina NovaSeq platform. Information on the sequencing data generated by Guangzhou Nuosai Biotechnology Co., Ltd. and used for further assembly and annotation is shown in Table 1. The paired-end read sequences of the four species were submitted to the National Center for Biotechnology Information (NCBI) database (BioProject ID: PRJNA931566).

### Assembly and Annotation of cpDNA

The cpDNA were extracted and assembled from the original data using GetOrganelle V1.7.5.3 software (Jin et al., 2020). After the cpDNA were assembled, the chloroplast genome loop with a typical tetrad structure was concatenated and visualized using Bandage software (Wick et al., 2015). The chloroplast genome of *C. vietnamennsis* (NC_060778.1) was used as the reference genome, and the sequenced genomes were annotated using PGA software (Qu et al., 2019). Geneious 9.0.2 software was used to visualize the annotated sequences (Kearse et al., 2012), and manual adjustments were made to improve the sequences. The annotation results were submitted to the National Center for Biotechnology Information (NCBI) under the accession numbers ON367462, ON418964, ON418963, and ON418965. The online software OGDRAW (Greiner et al., 2019) was used to map the cpDNA.

### Sequence Alignment Analysis of cpDNA

The four cpDNA were assembled and uploaded to NCBI. The cpDNA of *C. vietnamensis, C. crapnelliana, C. oleifera, C. chekiangoleosa, C. gauchowensis*, and *C. kissii* were downloaded to conduct analyses. The GenBank accession numbers were NC060778, KF753632, KY406750, NC037472, NC053541, and NC053915, respectively (**Supplementary table S1**). ShufflemVISTA software (https://genome.lbl.gov/vista/mvista/submit.shtml) was used to align the cpDNA of the different species using the alignment subprogram Shuffle-LAGAN with global pair-wise alignment of the finished sequences (Chris et al., 2000). The online software IRscope (https://irscope.shinyapps.io/irapp/) (Amiryousefi et al., 2018) was used to make comparisons of the annotated cpDNA of *C. semiserrata, C. meiocarpa, C. suaveolens*, and *C. osmantha* with the downloaded sequences in the inverted repeat (IR) regions of the cpDNA, including regions of contractions and expansions. The molecular markers of the cpDNA for the above 10 *Camellia* species were developed using DnaSP6 software (Rozas et al., 2017).

### Repeat Sequence Analysis and Codon Bias Analysis

The online software MISA (https://webblast.ipk-gatersleben.de/misa/) (Beier et al., 2017) was used to localize repetitive elements in the cpDNA of *C. semiserrata, C. meiocarpa, C. suaveolens, C. osmantha*, and *C. vietnamensis*. To detect simple sequence repeats (SSRs), the minimum number of repetitions for mononucleotide, dinucleotide, trinucleotide, tetranucleotide, and hexanucleotide repeats was set to 12, 6, 4, 3, 3, and 3, respectively. The online software REPuter (Kurtz et al., 2001) was used to identify interspersed nuclear elements. Four types of repeat sequences were detected, including forward repeat (F), reverse repeat, palindromic repeat (P), and complementary repeat sequences. The minimum repeat size was set at 30 bp, and the minimum repeat distance was set at 3, respectively. Statistical analyses were conducted using Microsoft Excel 2019, and figures were made using Origin 2022.

The protein-coding genes (CDS) of *C. semiserrata, C. meiocarpa, C. suaveolens*, and *C. osmantha* were extracted using Geneious 9.0.2. To reduce the errors caused by short sequences, duplicate sequences and sequences less than 300 bp in length were removed from these coding sequences. Next, coding sequences that start with an ATG and end with a TAA, TGA, and TAG were selected. Finally, CodonW 1.4.2 software was used to conduct relative synonymous codon usage (RSCU) analysis on each of the CDS of each species to characterize patterns of codon bias (Sharp and Li, 1987).

### Phylogenetic Analysis

The cpDNA of 29 *Camellia* species and two outgroup species (*Schima superba* and *Tutcheria pingpiensis*) were downloaded from the NCBI database to construct phylogenetic trees with the resequenced cpDNA of *C. semiserrata, C. meiocarpa, C. suaveolens*, and *C. osmantha* (**Supplementary Table S1**). Geneious 9.0.2 software was used to select sequences of regions from the large single-copy (LSC), IR, small single-copy (SSC), and 68 CDS. A multiple sequence alignment of the whole genome and the selected sequences was conducted using MAFFT v7.308 software (Kazutaka et al., 2017). A phylogenetic tree was constructed for each sequence. Phylogenetic trees were constructed using MrBayes ver. 3.2.7a(http://nbisweden.github.io/MrBayes/index.html) and RAxML software with 1, 000 bootstrap replicates (Ronquist et al., 2012; Stamatakis, 2014). FigTree 1.4.4 (http://tree.bio.ed.ac.uk/software/figtree/) was used to build the Bayesian and maximum likelihood (ML) tree, and the iTOL online tool (https://itol.embl.de/) was used to visualize the phylogenetic relationships (Letunic and Bork, 2021).

## Results

### Basic Characteristics of the cpDNA of Four Species

The cpDNA of the four *Camellia* species exhibited a typical tetrad structure (**Figure 1 and Table 2**), with a total length ranging from 156, 965 to 157, 018 bp, consisting of an LSC region (ranging from 866, 647 to 86, 656 bp), an SSC (ranging from 18, 282 to 18, 408 bp), and a pair of IR regions (ranging from 25, 954 to 26, 042 bp). The lengths of the four cpDNA only varied by 53 bp. The cpDNA of *C. semiserrata* was the longest, and that of *C. meiocarpa* was the shortest. The LSC region was the least variable region in the cpDNA of the four *Camellia* species (only varying by 9 bp), whereas the SSC region was the most variable region among the four *Camellia* species (varying by 126 bp). The average GC content of the four chloroplast genomes was 37.3%. The GC content of the LSC region was 35.3%, and that of the SSC region ranged from 30.5 to 30.6%. The GC content of the IR region (43.0%) was higher than that of the LSC and SSC regions. A total of 134 genes were identified in the four cpDNA, including 88 CDS, 37 transfer RNA (tRNA) genes, and 8 messenger RNA (mRNA) genes. The pseudogene ycf1 was located in the boundary region between the IRB and SSC regions, and other incomplete copy of *ycf1* was also located at the boundary between the SSC and IRA regions. The cpDNA of the four *Camellia* species are highly conserved given that they all possess the basic structural features of cpDNA.

**Table 2.**
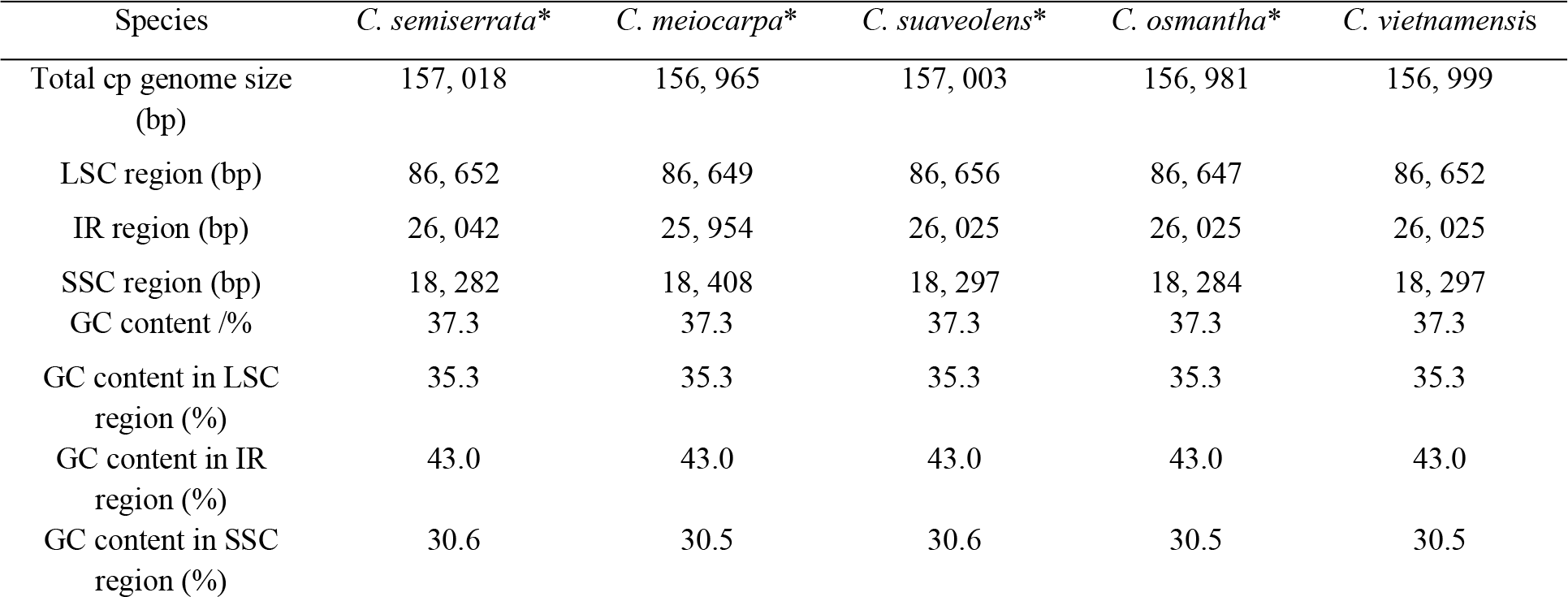
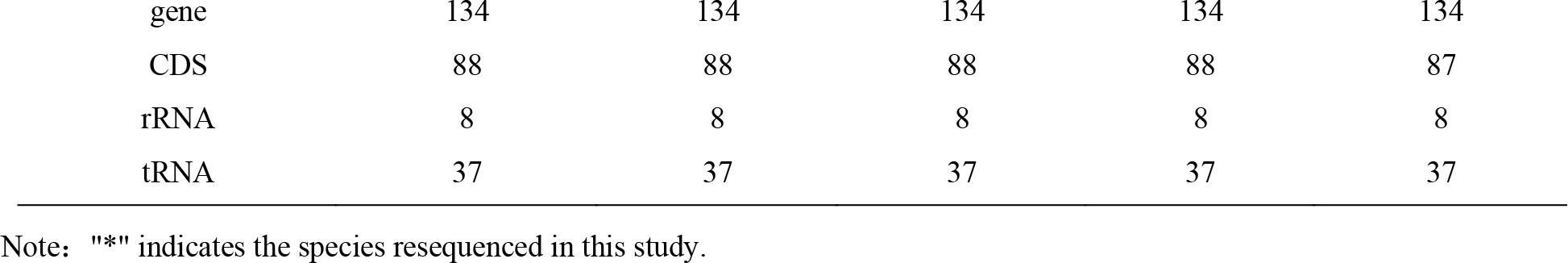
Analysis of the cpDNA characteristics of four *Camellia* species.

**Figure 1.**
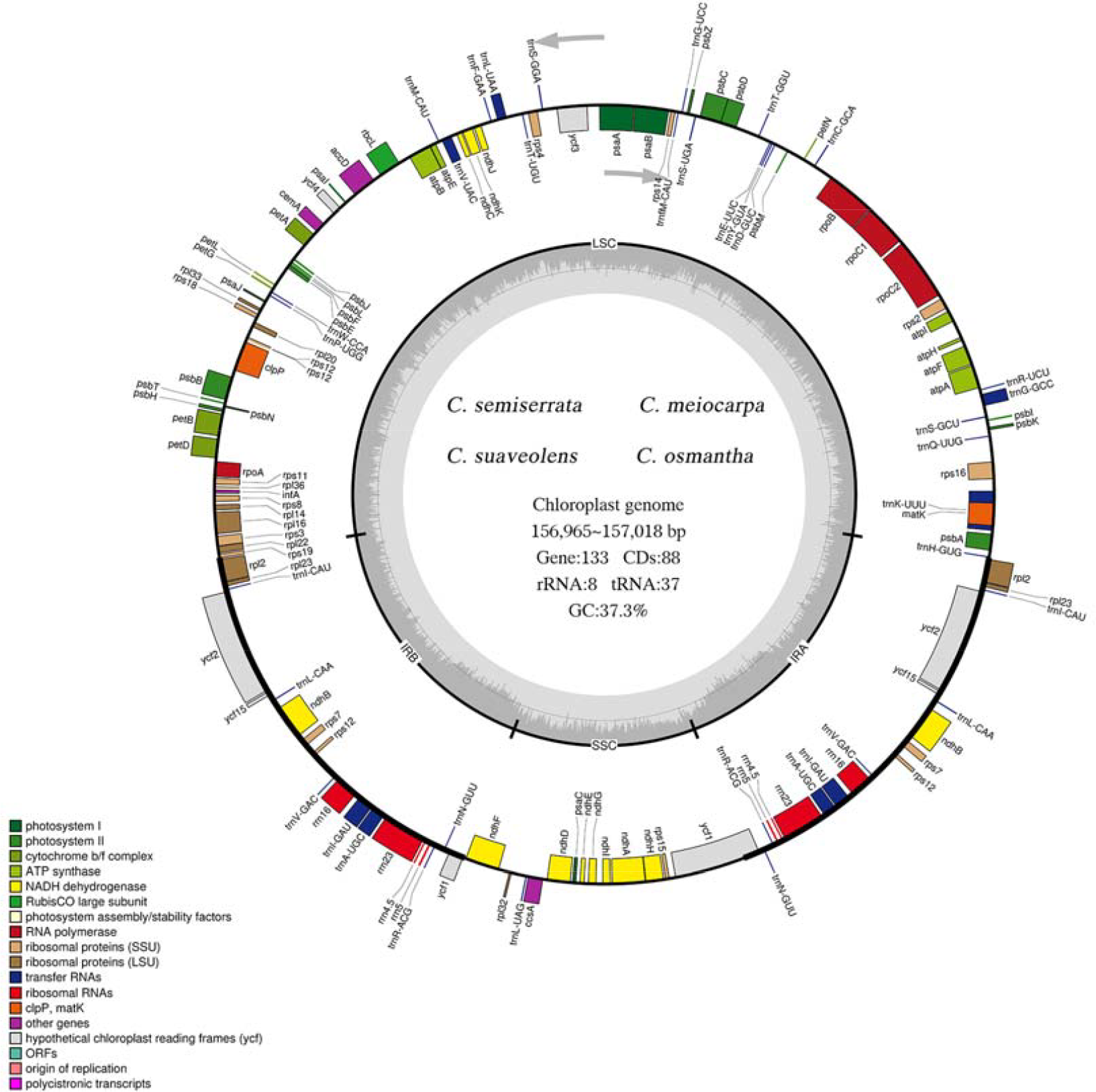
Gene map of the cpDNA of four *Camellia* species. The genes inside the circle are transcribed clockwise, and the genes outside is transcribed counterclockwise. Genes with different functions are color-coded. The darker gray in the center circle indicates the GC content, and the lighter gray indicates the AT content.

Excluding the one pseudogene, 98 out of 133 genes were unique, including 75 CDS and 23 tRNA genes. The other 35 genes were located in the IR regions, including 13 CDS (*ycf1, rps7* *2, *ndhB*\*2, *ycf15* *2, *ycf2* *2, *rpl23* *2, and *rpl2* *2), 8 ribosomal RNA (rRNA) genes, and 14 tRNA genes. A total of 45 genes were involved in photosynthesis, 75 genes were involved in self-replication, 6 genes encoded other proteins, and 7 genes had unknown functions. Sixteen genes had one intron, and two genes had two introns (**Table 3**).

**Table 3.**
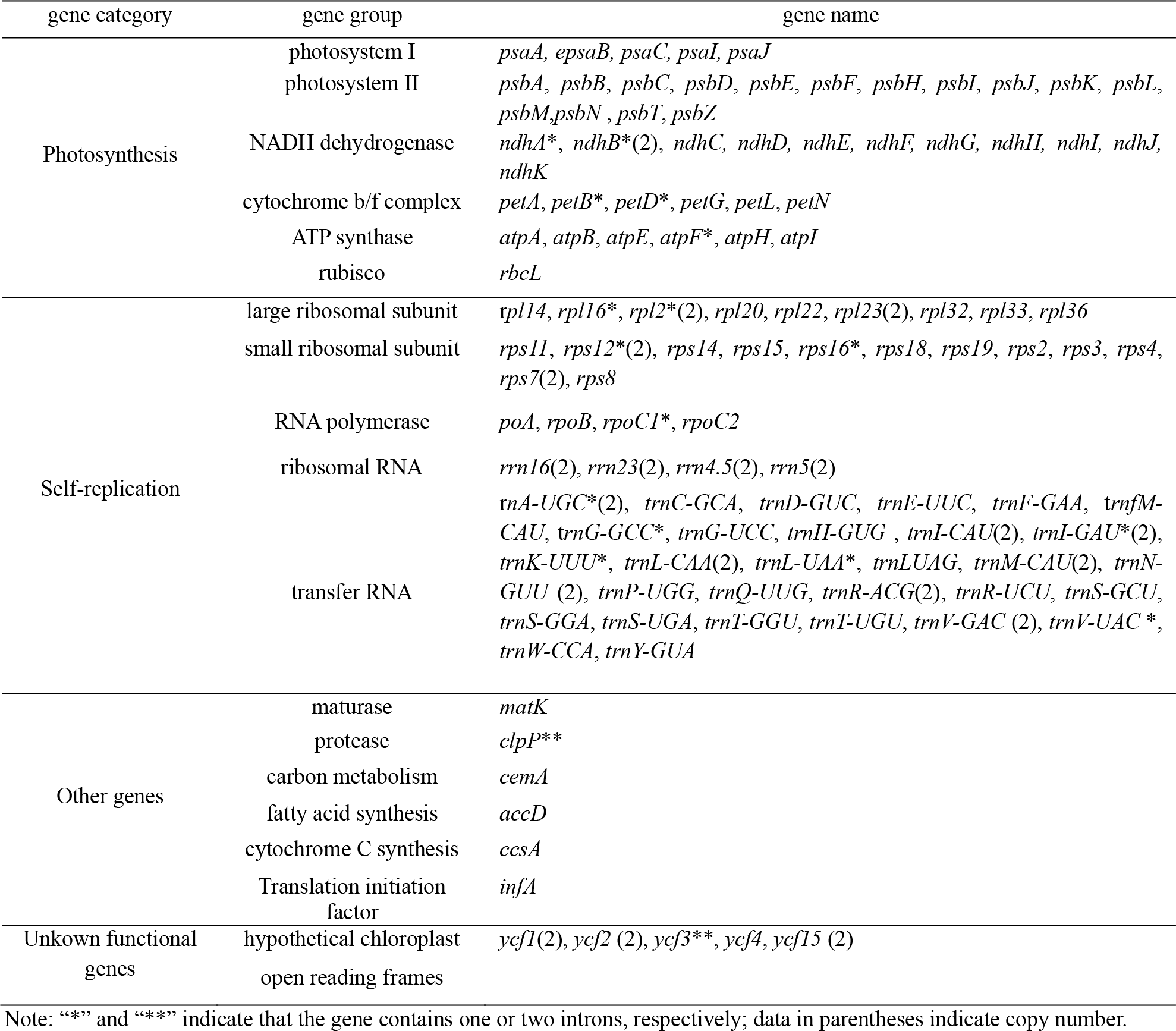
List of genes found in the cpDNA of four *Camellia* species.

### Contractions and Expansions of the IR Boundary

IR boundary analysis of 10 *Camellia* species, including the four *Camellia* species in this study (**Figure 2**), revealed that the structure and sequences of the IR boundary regions of *C. semiserrata, C. meiocarpa, C. suaveolens, C. osmantha*, and *C. vietnamensis* were similar. The genes of *rpl22, rps19, rpl2, ycf1, ndhF*, and *trnH* were mainly located near the IR/LSC and IR/SSC boundaries of the cpDNA for these 10 *Camellia* species. The *rps19* gene crossed the LSC/IRB boundary in nine of these species. The IR region of *C. chekiangoleosa* has undergone a contraction. Consequently, the *rpl2* gene has expanded to the LSC region and crosses the LSC/IRB boundary. The *rpl2* gene in *C. kissii* and *C. gauchowensis* was not present in the LSC/IRB boundary. In *C. semiserrata, C. meiocarpa, C. suaveolens, C.osmantha*, and *C. vietnamensis, ycf1* is a pseudogene in the SSC/IRA boundary, and there is also an incomplete copy of *ycf1* in the SSC/IRB boundary region. The *ycf1* gene in *C. meiocarpa* is located 1 bp away from the SSC/IRA boundary. And the *ycf1* gene in the other four species is located 106 bp away from the SSC/IRA boundary. The locations of *ndhF* and *ycf1* in the SSC/IRB and SSC/IRA boundary regions vary in *C. kissii, C. oleifera, C. crapnelliana*, and *C. chekiangoleosa*. This is mainly associated with the contraction and expansion of the IR and SSC regions. The locations of *rpl2* and *trnH* in the LSC/IRA boundary region are consistent in other plants, with the exception of *C. gauchowensis* and *C. crapnelliana*, which did not possess the *rpl2* gene.

**Figure 2.**
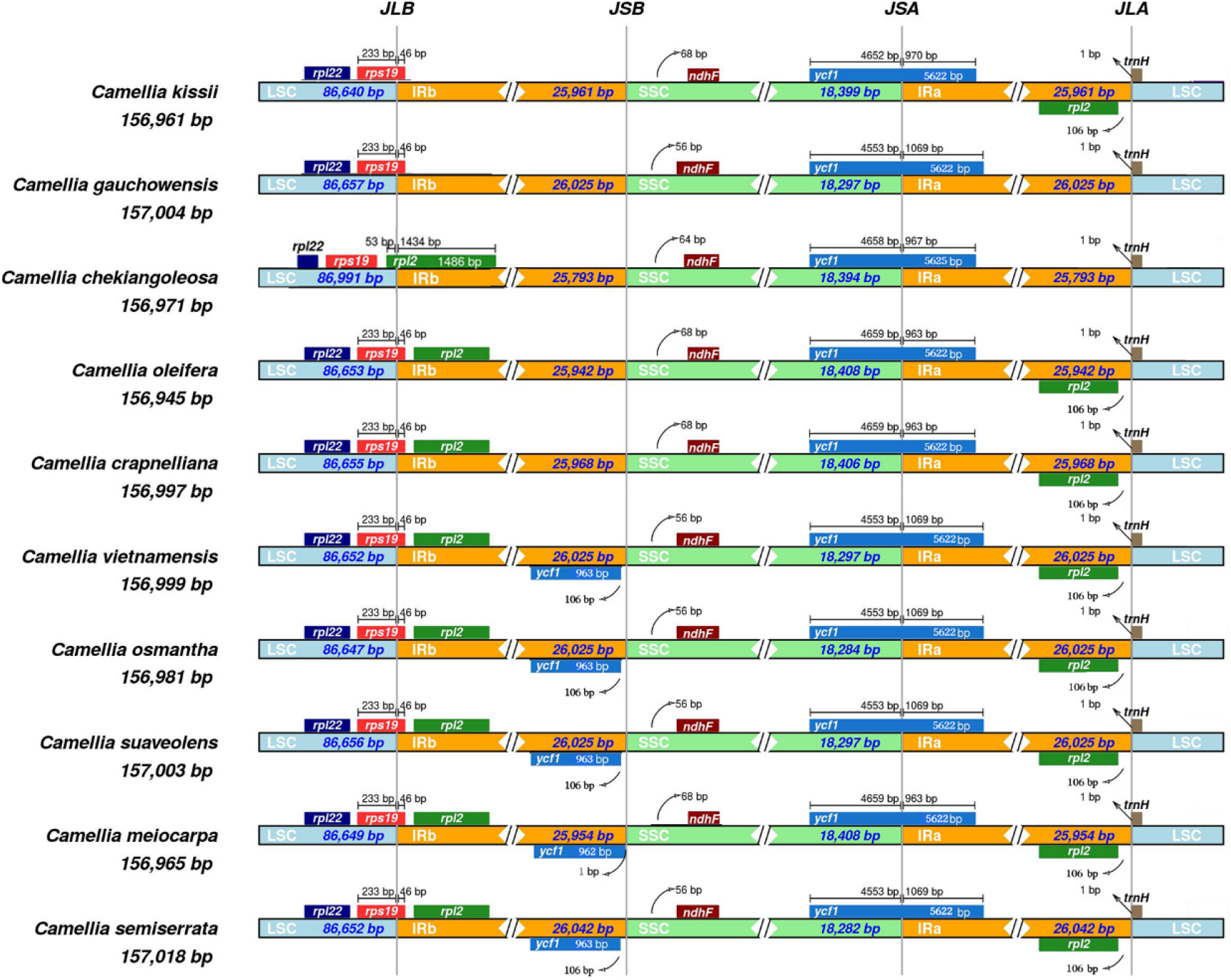
Comparison of IR/SC boundary regions of 10 *Camellia* species. LSC, Large single-copy; SSC, Small single-copy; IRA and IRB, inverted repeats. JLB, junction between LSC and IRB; JSB, junction between SSC and IRB; JSA, junction between SSC and IRA; JLA junction between LSC and IRA.

### Molecular Marker Detection

The cpDNA of 10 *Camellia* species, including the four focal *Camellia* species in this study, were compared with mVISTA software using *C. vietnamensis* as a reference (**Figure 3**). The results showed that sequence similarity in the coding region was high. However, sequence similarity of the non-coding region was low. The LSC region was the most variable region across the cpDNA, followed by the SSC, IRB, and IRA regions. Thus, variation in the IR region is low, which suggests that it is evolutionarily conserved. The most conserved genes were rRNA genes, as no significant variation was observed in rRNA genes among cpDNA.

**Figure 3.**
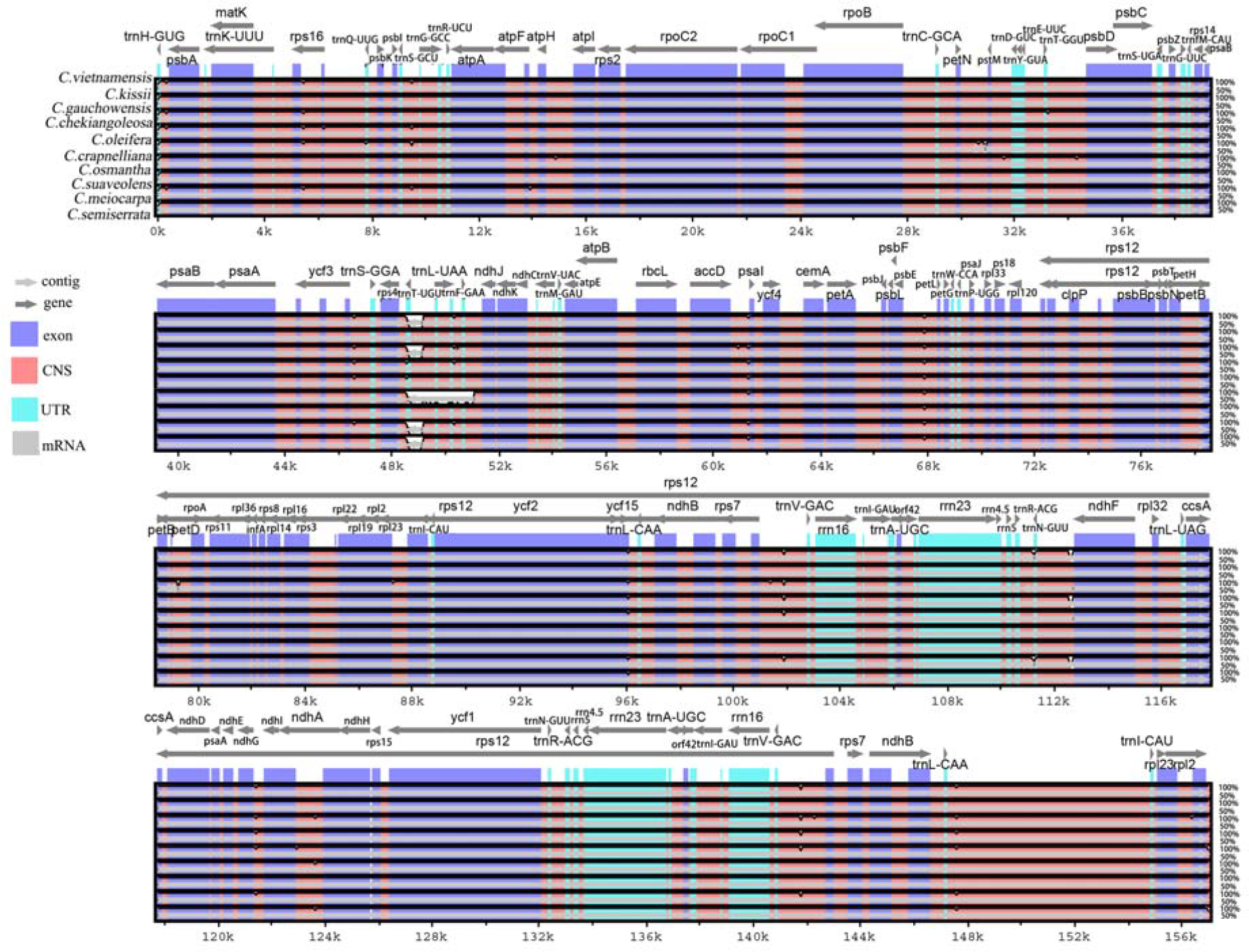
Comparison of the cpDNA of 10 *Camellia* species. The X-axis indicates aligned base sequences, and the Y-axis indicates percent pairwise identity within 50-100%. Grey arrows represent genes and their directions. Blue boxes indicate exon regions, light blue boxes indicate regions encoding RNA genes, and red boxes indicate non-coding sequences.

Molecular markers were developed for these 10 *Camellia* species (**Figure 4**). Five highly variable gene spacers or genes were identified, including *trnQ-UUG*—*trnG-GCC, petN*-*pstM*, and *trnL-UAA*—*ndhJ* in the LSC region, *ycf15*—*ndhB* in the IR region, and *ycf1* in the SSC region. In the *trnQ-UUG*—*trnG-GCC* region from the LSC region, the pi value was highest from 9, 296 to 9, 905 bp (0.01737). This region has a total of 21 mutated sites and, GC content of 30.7%. It was thus the most highly mutated region in the entire cpDNA. In the *ycf15*—*ndhB* region of the IR region, pi was highest in the 96,233–96,837-bp, 96,438–97,037-bp, and 147,251–147,850-bp regions, all of which were approximately 0.00437. There were total of six mutated sites, and the GC content was 42.8% in this region. In the ycf1 gene in the SSC region, pi was highest in the 128,406–129,005-bp region (0.00463) with nine mutated sites, and a GC content of 23.7%. These regions with high variability can be used as DNA barcodes for species identification.

**Figure 4.**
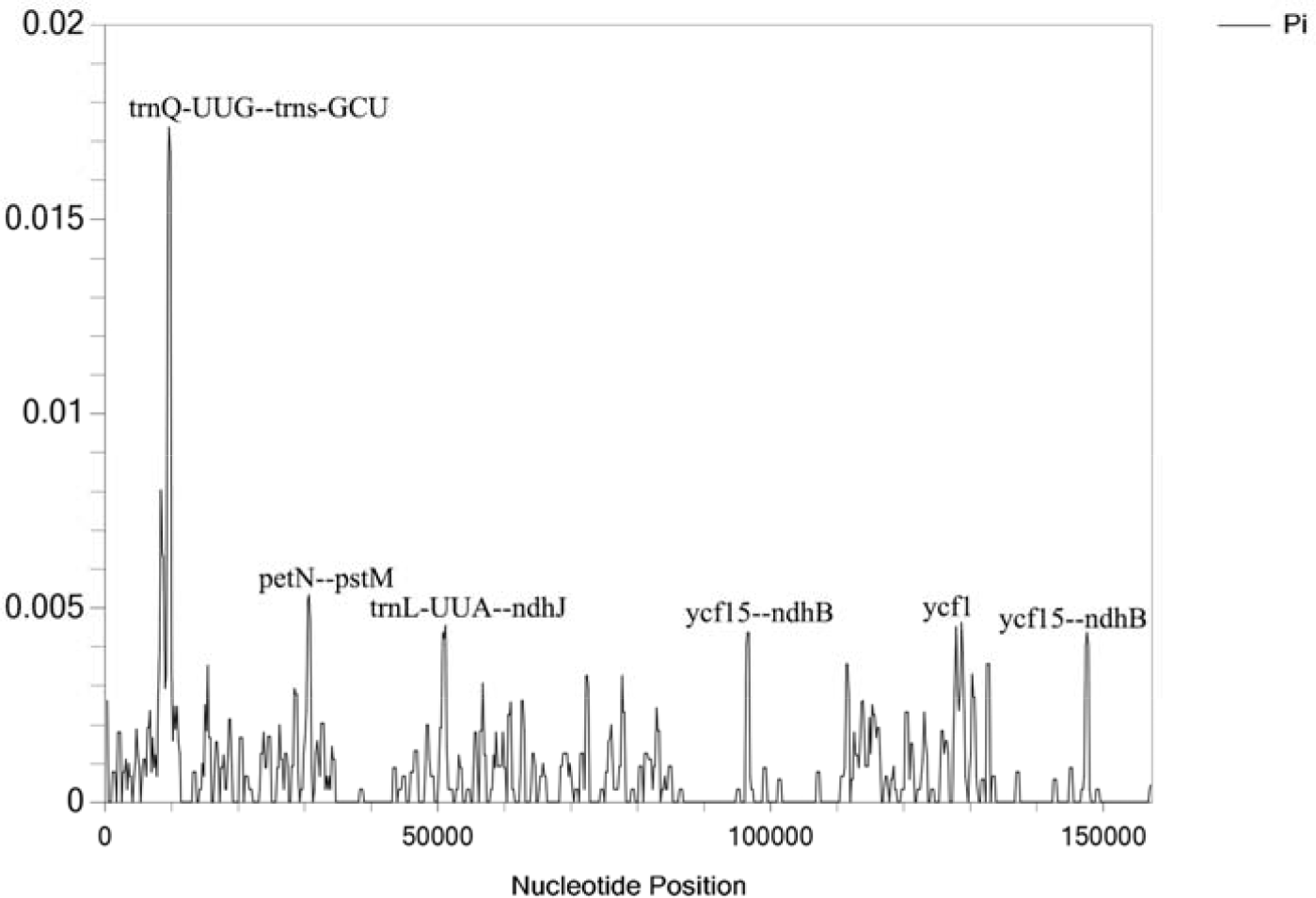
Chloroplast genome sliding window analysis of 10 *Camellia* species. Window length: 2000 bp; step size: 200 bp. X-axis: the position of the midpoint of a window. Y-axis: nucleotide diversity of each window.

### Codon Bias Analysis

A total of 88 protein-coding genes (CDS) were identified in the cpDNA of *C. semiserrata, C. meiocarpa, C. suaveolens*, and *C. osmantha* using Geneious 9.0.2 software. The lengths of the CDS were greater than 300 bp, and the start codon of these CDS was ATG. The stop codons were TAA, TGA, and TAG. Totally of 51 sequences with each kind of stop codon were obtained. There were total of 20,747, 20,743, 20,743, and 20,780 codons in the 51 CDS from *C. semiserrata, C. meiocarpa, C. suaveolens*, and *C. osmantha*, and the GC content of their CDS was 37.47%, 37.46%, 37.45%, and 37.45%, respectively (**Supplementary Table S2**). RSCU analysis of these codons revealed (**Figure 5** and **Supplementary Table S2**) high conservation in the codon usage of the four *Camellia* plants. The highest coding rate was observed for the codon UUA, which codes for leucine, and UUA comprised 698, 697, 698, and 699 codons in *C. semiserrata, C. meiocarpa, C. suaveolens*, and *C. osmantha*, respectively. The RSCU value of UUA was 1.97. The lowest coding rate in *C. semiserrata, C. meiocarpa*, and *C. suaveolens* was observed for AGC (84), which codes for serine (Ser), and its RSCU value was 0.32. The coding rate of AGC and CGC (85 and 69, respectively), which codes for arginine (Arg), was lowest in C. osmantha. The RSCU value for CGC was 0.33. The RSCU value was greater than one for 29 codons across the four *Camellia* species, with the exception of stop codons. 12 of these codons ended in A, 16 ended in U, and one ended in G. Two codons, AUG and UGG, had RSCU values close to 1, indicating the absence of codon usage bias for these two codons. The RSCU value was less than 1 for 30 codons, with the exception of stop codons. Number of 16 of these codons ended in C, 12 ended in G, and 2 ended in A. These findings indicate that there is a bias for codons ending with A or U in the cpDNA of *C. semiserrata, C. meiocarpa, C. suaveolens*, and *C. osmantha*.

**Figure 5.**
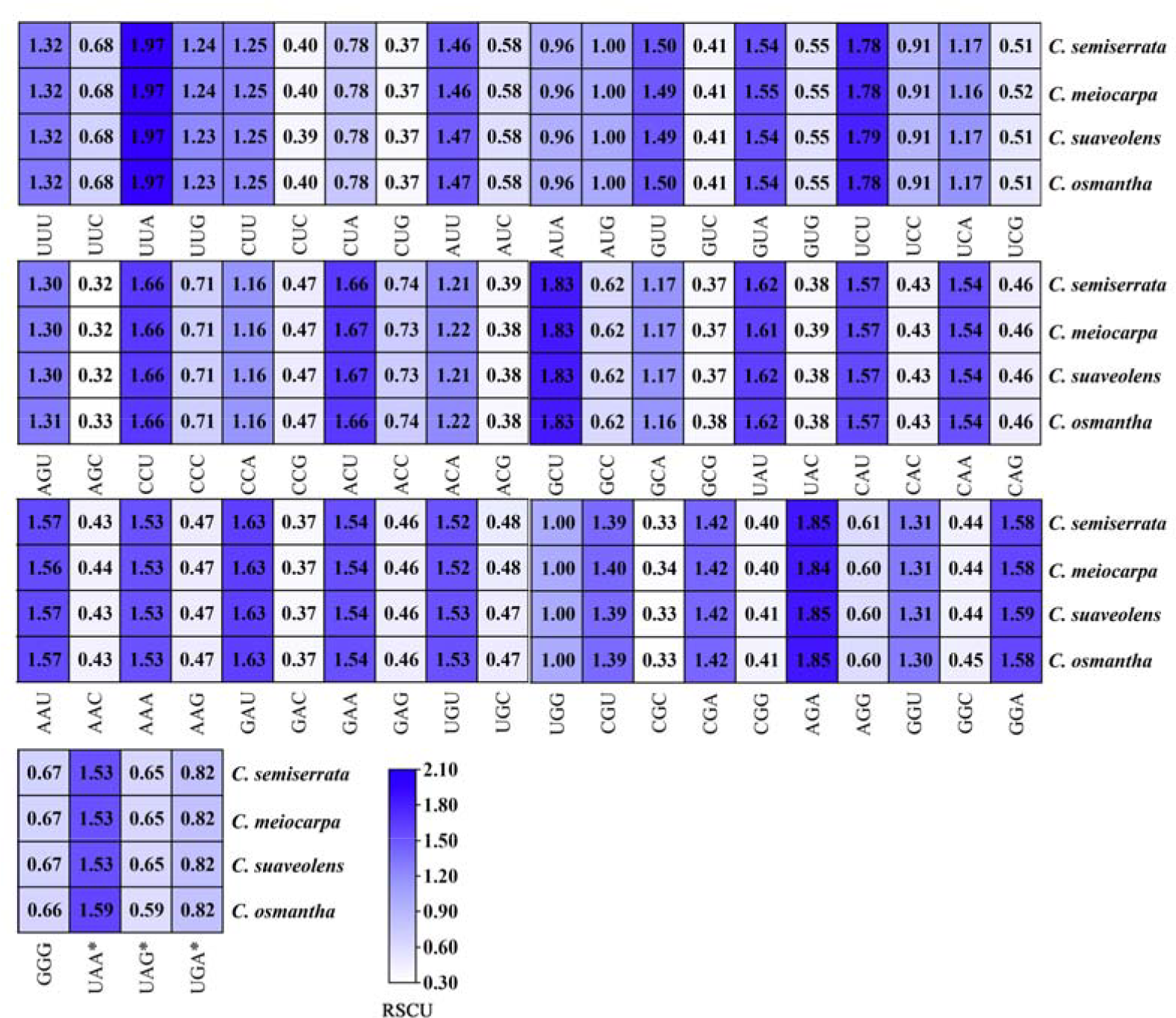
RSCU analysis of the cpDNA of four *Camellia* species. Darker colors indicate, greater RSCU and skew. “*” indicates the termination codon.

### SSR and Repeat Sequence Analysis

SSR analysis of the cpDNA of four *Camellia* plants and six other plant species was conducted. Dinucleotide and pentanucleotide repeats were not detected in the 10 *Camellia* species (**Figure 6A**). The number of SSRs ranged from 36 to 45, and the greatest number of SSRs was observed in *C. crapnelliana*. The lowest number of SSRs was observed in *C. chekiangoleosa*. Mononucleotide repeats were the most common, followed by tetranucleotide, trinucleotide, and hexanucleotide repeats. The numbers of SSRs in the four *Camellia* species ranged from 38 (in *C. meiocarpa*) to 40 (in *C. osmantha*). Hexanucleotide repeats were not detected in *C. semiserrata* and *C. suaveolens*. The number of mononucleotide and trinucleotide repeats in *C. meiocarpa* was 22 and one, lower than *C. osmantha, C. semiserrata* and *C. suaveolens*. The number of SSRs was lowest in *C. meiocarpa* among the four species.

**Figure 6.**
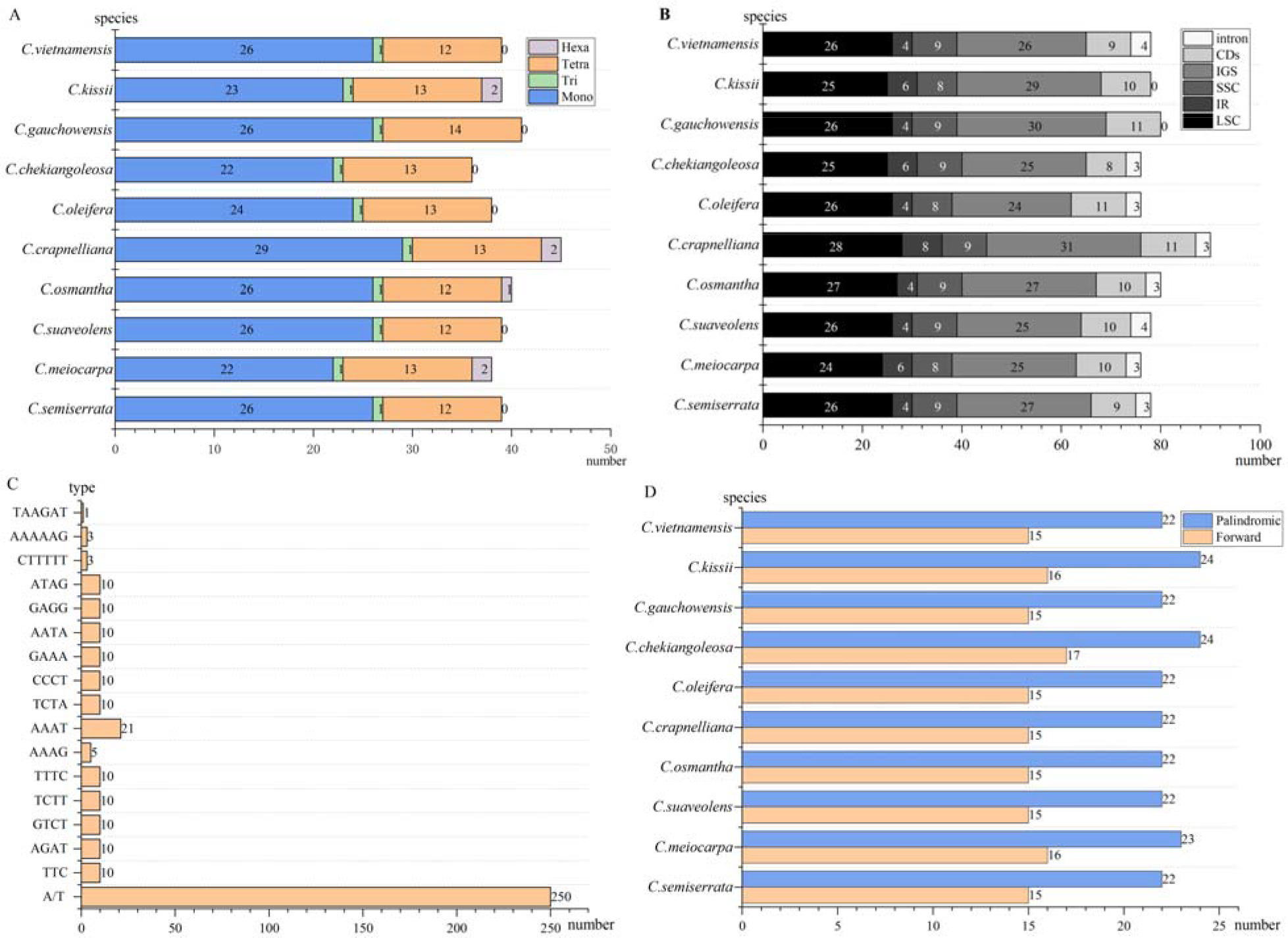
SSRs and INE analysis of the cpDNA of 10 *Camellia* species. X-axis: the number of SSRs or INE; Y-axis: species or SSR type. (A) Number of SSRs; (B) distribution of SSRs; (C) type of SSRs; and (D) type and number of INEs.

SSRs in the four *Camellia* species were most abundant in the LSC region and least abundant in the IR region (**Figure 6B** and **Figure 6C**). There were two main types of mononucleotide repeats (A/T), and these were only distributed in the LSC and SSC regions. The number of mononucleotide repeats was greater in the LSC region than in the SSC region. Only one type of trinucleotide repeat (TTC) was observed, and it was only present in the LSC. There were 12 types of tetronucleotide repeats across all partitions (AGAT/GTCT/TCTT/TTTC/AAAT/AAAG/TCTA/CCCT/GAAA/ AATA/GAGG/ATAG). The most common tetranucleotide repeat was the AAAT type, and the least common was AAAG. Tetranucleotide repeats were most common in the LSC region, followed by the SSC and IR regions. AAAG was only observed in *C. meiocarpa*. Three hexanucleotide repeats were observed with CTTTTT/AAAAAG/TAAGAT) distributing in the IR region only. These hexanucleotide repeats were most abundant in the gene spacer region and least abundant in introns.

The results of the repeat sequence analysis are shown in **Figure 6D**. The number of repeat sequences in the cpDNA of these 10 *Camellia* species ranged from 37 to 41, and the lengths of these sequences ranged from 30 to 64 excluding the IR region. The palindromic (P) sequences identified were always more abundant than the forward (F) sequences. A total of 37 repeats were detected in *C. semiserrata, C. suaveolens*, and *C. osmantha*, including 15 positive repeats and 22 P sequences. There were 39 repeat sequences, 16 F sequences, and 23 P sequences in *C. meiocarpa*.

### Phylogenetic Analysis

The aligned sequences of the complete cpDNA (**Figure 7A**), LSC region (**Figure 7B**), SSC region (**Figure 7C**), IR regions (**Figure 7D**), and shared CDS (**Figure 7E**) of the 35 species were used to construct phylogenetic trees via the Bayesian and ML methods, respectively. Most of the relationships in the phylogenies built based on the complete cpDNA and LSC region had moderate to high support, and the general topology of the trees based on the LSC region and the complete cpDNA was similar (**Figure 7**). In this phylogenetic tree, *C. semiserrata, C. osmantha*, and *C. suaveolens* were nested within their own small clades that were clustered into a large clade with *C. meiocarpa*. The phylogenetic trees generated from the first four datasets showed that *C. semiserrata* was most closely related to *C. alzalea*, while *C. suaveolens* was most closely related to *C. gauchowensis. C. osmantha* was most closely related to *C. vietnamensis* and was similar to *C. suaveolens* and *C. gauchowensis*. Species of *C. meiocarpa* was most closely related to *C. oleifera*. Phylogenetic trees of *Camellia* species based on the IR regions, the SSC region, and the shared CDS were not highly supported. Thus, the IR regions, the SSC region, and CDS are more conserved and less variable.

**Figure 7.**
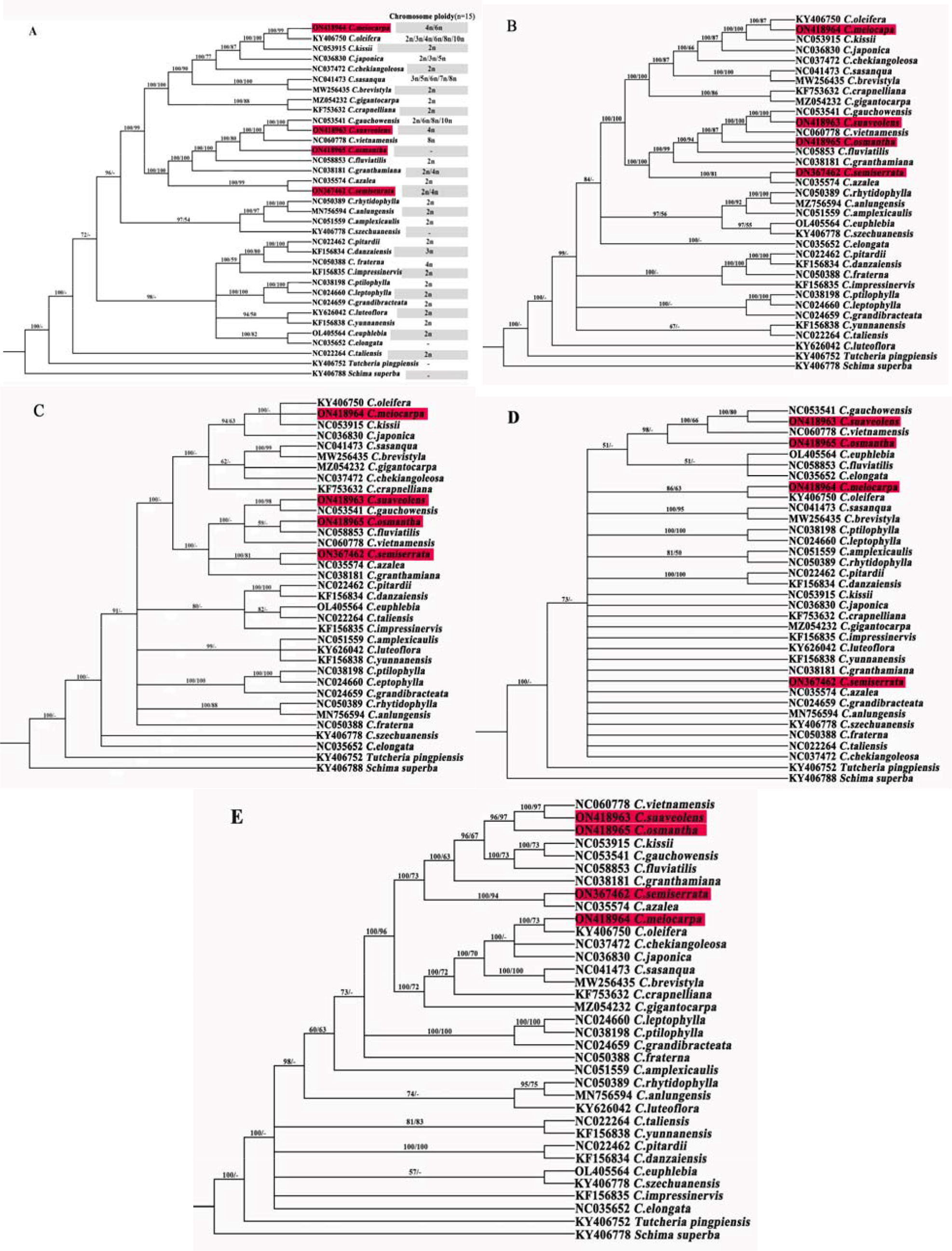
Bayesian phylogenetic tree based on 35 species of complete cpDNA, LSC, SCC, IR and CDS. (A) Bayesian phylogenetic tree based on the complete cpDNA; (B) Bayesian phylogenetic tree based on the LSC region; (C) Bayesian phylogenetic tree based on SSC regions; (D) Bayesian phylogenetic tree based on the IR regions; and (E) the Bayesian phylogenetic tree based on the CDS. The numbers on the branches are Bayesian and ML bootstrap values.

## Discussion

The structure, size, gene content, and genotypes of the cpDNA for most land plants are highly conserved. The lengths of the cpDNA of 100 species in the family Theaceae in the NCBI database range from 150 to 160 kb, and all of them have a typical tetrad structure. The degree of sequence conservation is positively correlated with the GC content (Zheng et al., 2022). The cpDNA of *C. semiserrata, C. meiocarpa, C. suaveolens*, and *C. osmantha* are highly conserved in their structure, size, gene content, and gene type. The resequenced complete cpDNA of these four *Camellia* species provide new genomic resources for *Camellia* in the NCBI database (Dong et al., 2021; Tong et al., 2022; Chen et al., 2022). The IR region is considered the most conserved region in the cpDNA, and contractions and expansions of the IR boundary play important roles in determining the size and evolutionary trajectory of cpDNA (Wang et al., 2018). In most higher plants, the evolutionary rate of the cpDNA is relatively low, while the sequences and structure are highly conserved. However, the contractions and expansions of IR boundary are commonly observed (Zheng et al., 2022). Contractions and expansions of the IR regions in the four focal *Camellia* species in this study were similar to those observed in *C. weiningensis* (Li et al., 2020), *C. tienii* (Ding et al., 2022), and *C. japonica* (Li et al., 2019). These findings indicate that patterns of IR contractions and expansions vary among *Camellia* species, and this results in changes in the relative lengths of tetrads and full-length cpDNA.

Analysis of nucleotide repeats and molecular makers in cpDNA can be used to distinguish between populations or species, and provide insights into patterns of genetic diversity and systematic relationships (Tang et al., 2022; Guisinger et al., 2010). Highly variable loci in chloroplasts can provide substantial DNA barcoding information that can be used to identify plants. Analysis of the *trnl-trnF, rpl16*, and *psbA*-*tmH* sequences of Sect. Chrysantha in *Camellia* species has shown that *rpl16* and *psbA*-*tmH* sequences can be used to accurately identify *C. pubipetala* (Chen et al., 2021). The *trnH*-*psbA* sequence can be used to distinguish plants in different *Camellia* species, but its effectiveness for interspecific classification is weak (Wen et al., 2017). In our study, five highly variable regions were detected (*trnQ-UUG*—*trnG-GCC, petN*-*pstM, trnL*-*UAA*—*ndhJ, ycf15*—*ndhB*, and *ycf1*), and these regions could be used in future taxonomic studies of oil-seed *Camellia* species. SSR markers have been widely used to analyze the phylogenetic relationships and genetic diversity of *Camellia* species (Dong et al., 2022; Tao et al., 2019; Wang, 2019; Zhang et al., 2018). Variation was observed in the number of SSRs in the four *Camellia* species and ranged from 36 to 45. The dinucleotide repeats and pentanucleotide repeats were not detected in these four *Camellia* species. No hexanucleotide repeats were detected in *C. semiserrata* and *C. suaveolens*. The numbers of different types of SSRs among the *Camellia* species in this study differ from those of other *Camellia* species in previous studies (Yin et al., 2018). The results of our study showed that SSRs were most common in non-coding regions, followed by regions of coding, LSC, SSC, and IR. The most common mononucleotide repeats were A and T, and no pentanucleotide repeats were observed, which consistent with the results of previous studies (Ding et al., 2022; Yin et al., 2018; Yang et al., 2013). The SSRs of the cpDNA in four *Camellia* species comprised 37 to 39 repeats with lengths ranging from 30 and 64 bp, which is consistent with the results of a previous study of *C. tienii* (Ding et al., 2022). In this study, we did not identify genes or molecular makers in cpDNA that mediated increases in the oil deposited in the seeds as has been identified in other species, such as the long-chain acyl-coA synthetase gene in the FA and TAG synthesis pathways in the leaves of *Suaeda salsa* (Gao et al., 2018).

Codon bias affects mRNA stability, mRNA transcription, and the accuracy of protein translation and protein folding and thus plays a key role in regulating gene expression (Ren et al., 2019). Information on the codon usage of the cpDNA in *Camellia* species, especially comparisons among species, can provide insights into differentially expressed genes, optimize codon usage, and aid the selection of varieties with desirable characters (Teng et al., 2021; Lai et al., 2022; Zhou et al., 2022). The codons of the cpDNA in the four *Camellia* species mainly ended in A/U. This is consistent with the observed codon bias in *C. nitidissima, C. oleifera*, and *C. osmantha* in previous studies (Wang et al., 2018; Geng et al., 2022; Hao et al., 2022).

Frequent interspecific hybridization and polyploidy have hindered efforts to resolve the phylogeny and taxonomy of the genus *Camellia*. Comparative analysis of whole cpDNA can provide more reliable insights into phylogenetic relationships among *Camellia* members (Li et al., 2019; Jiang, 2017). Phylogenetic trees built based on the IR dataset, SSC dataset, and CDS were not highly supported, indicating that data from these regions of the cpDNA should not be used in phylogenetic studies of *Camellia* species. The topologies of phylogenetic trees based on the complete cpDNA dataset and the LSC dataset were highly supported (**Figure 7A**). The phylogenetic trees obtained in this study revealed that (1) *C. semiserrata* was most closely related to *C. azalea*; (2) *C. suaveolens* was most closely related to *C. gauchowensis*; and (3) *C. osmantha* was most closely related to *C. vietnamensis*, which was similar to *C. suaveolens* and *C. gauchowensis*.

C. *semiserrata* is more similar to *C. japonica* and *C. chekiangoleosa* based on cytological and morphological characters (Ni, 2007), yet this finding is inconsistent with the results of our study. However, a study of interspecific hybridization of *C. azalea* and *C. semiserrata* has revealed high hybridization affinity among these species (Zhong et al., 2020), and all three of these species are diploid (Jia, 2015). *C. gauchowensis* was considered a synonym of *C. vietnamensis* in Flora of China. However, *C. suaveolens* was more closely related to *C. gauchowensis* than to *C. vietnamensis* according to phylogenetic trees based on cpDNA in this study (**Figure 7A)**. The cross-compatibility between *C. gauchowensis* and *C. suaveolens* via reverse hybridization is low because of their different ploidy. *C. vietnamensis* and *C. gauchowensis* show high cross-compatibility in reciprocal crosses with the same chromosome ploidy (Zhang et al., 2022). The results of ploidy and hybridization analysis revealed that *C. gauchowensis* is more closely related to *C. vietnamensis* than to *C. suaveolens. C. suaveolens* has recently been shown to be more similar to *C. furfuracea, C. osmantha*, and *C. fluviatilis* based on morphological data and inter-simple sequence repeat markers (Jia, 2015; Wang et al., 2004; Liang et al., 2017). This finding is also not consistent with the results of this study. *C. suaveolens* and *C. gauchowensis* are both decaploid (Zhang et al., 2022). *C. osmantha* shows high affinity with *C. gauchowensis* when hybrids are used as the male parent (Ma et al., 2012). In our study, *C. osmantha* was more closely related to *C. vietnamensis* than to *C. gauchowensis* and *C. suaveolens*. We also found that *C. meiocarpa* and *C. oleifera* were closely related. These findings are consistent with the current classification of these taxa (Jiang, 2017; Ni, 2007). *C. meiocarpa* and *C. oleifera* have been documented to hybridize. *C. meiocarpa* and *C. oleifera* were shown to be genetically closely related via unweighted pair group method with arithmetic mean clustering, and clear gene introgression associated with hybridization between these two species has been observed (Jia, 2015; Zhang et al., 2022; Huang, 2013). According to chromosome ploidy analyses, *C. meiocarpa* is hexaploid (Zhang et al., 2022), and *C. oleifera* ranged in diploid, tetraploid, hexaploid, or octaploid (Huang, 2013; Ye et al., 2020). Polyploidy is the result of interspecific hybridization (Liu and Huang, 2009). The gene exchange associated with interspecific hybridization between *C. meiocarpa* and *C. oleifera* might explain their high genetic similarity (Huang, 2013) and provides support for the taxonomic classification of *C. meiocarpa* and *C. oleifera* proposed by Zhang (1981). Most researchers follow the classification proposed by Hu Xianmu, in which *C. meiocarpa* is considered an independent *Camellia* species (Huang, 2013; Hu, 1957).

Sect. *Oleifera*, Sect. *Camellia*, Sect. *Paracamellia*, and Sect. *Furfuracea* did not form obvious clades in our trees, and many of the branches within these groups were not highly supported. The taxonomic classification of *Camellia* according to cpDNA data differs greatly from that based on morphological data (Lin et al., 2008; Shen et al., 2008; Zhang et al., 2016; Ye, 1988). This indicates that there are limitations associated with phylogenetic studies of *Camellia* based on cpDNA data. First, frequent hybridization and polyploidy of *Camellia* hinder its classification. Second, the cpDNA is uniparentally inherited, and it is evolutionarily conserved. Sequence differences between species are small, and the number of informative loci in the genome is not sufficiently high to permit the resolution of phylogenetic relationships among closely related taxa (Liu et al., 2015).

The cpDNA in this study, along with the results of hybridization and chromosome ploidy analyses, provided new insights into the evolutionary relationships among *Camellia* species, and this phylogeny is more robust than those constructed based on single genes. The general topology of the cpDNA tree is consistent with the classification of *Camellia* based on phenotypic data in this study (Wu et al., 2022). The publication of more nuclear genome sequences and the use of more information from natural hybridization and chromosome structure variation will likely provide further insights into the phylogenetic relationships among *Camellia* species.

## Conclusions

In this research, the cpDNA of four *Camellia* species, *C. semiserrata, C. meiocarpa, C. suaveolens*, and *C. osmantha*, were resequenced, assembled and annotated. We then analyzed the structure of the cpDNA of these four *Camellia* species, as well as contractions and expansions of the IR boundary, nucleotide polymorphism, repeat sequences and SSRs, codon bias, and clarified the phylogenetic relationships within *Camellia* based on hybridization and chromosome information. Our data will aid future studies of the identification, phylogenetic relationships, breeding, and sustainable development of germplasm resources of *Camellia* plants.

## Supplementary Material

Supplementary Table S1: Phylogenetic analysis of 35 species;

Supplementary Table S2: RSCU analysis of amino acids in the cpDNA of four *Camellia* species;

Supplementary_data_S1: The aligned sequences of the 35 cpDNA;

Supplementary_data_S2: The aligned sequences of the 35 CDS;

Supplementary_data_S3: The aligned sequences of the 35 IRB region;

Supplementary_data_S4: The aligned sequences of the 35 LSC region;

Supplementary_data_S5: The aligned sequences of the 35 SSC region.

## Data Availability Statement

The read sequences of *C. semiserrata, C. suaveolens, C. meiocarpa* and *C. osmantha* in this study were submitted to the NCBI database under the BioProject ID PRJNA931566 (https://www.ncbi.nlm.nih.gov/sra/PRJNA931566) and BioSample accessions numbers SAMN33062126, SAMN33062127, SAMN33062128, and SAMN33062129. The GenBank accession numbers were as follows: ON367462, ON418963, ON418964, and ON418965.

The name of the plant species *C. osmantha* is referred to as ‘Camellia sp. XJ-2021’ in the NCBI database under the GenBank accession number ON418963. The sequence information for *C osmantha* was published by Ye CX, Ma JL & Ye H. in Nanning, Guangxi, China, in 2012 (Ma et al., 2012). The information for ON418963 was uploaded to the NCBI database.

### Acknowledgements

We thank Xiaokeng Forest Farm (Shaoguan City, Guangdong Province, China) for providing plant materials. We thank Guangzhou Nuosai Biotechnology Co., Ltd. for helping us complete chloroplast genome sequencing.

